# Transient Room Lighting for Ambient Light Multiphoton Microscopy

**DOI:** 10.1101/2020.08.04.236364

**Authors:** A. Velten, A.J. Uselmann, S. Prajapati, J.S. Bredfeldt, T.R. Mackie, K.W. Eliceiri

## Abstract

Laser scanning microscopy techniques such as confocal and multiphoton fluorescence microscopy have been widely adopted by the biological research community due to their ability to monitor intact specimens at high spatial and temporal resolution. However, they have been limited for many biomedical, clinical and industrial applications by their fundamental need to operate in near absolute darkness. We present a lighting system that allows the use of light-sensitive imaging techniques in a fully-lit room by interleaving capture and illumination at a high frequency and exploiting the light averaging properties of the human eye. We use this system with a multiphoton fluorescence microscope to illustrate that this method is capable of image capture in a well-lit room on par with capture in absolute darkness. This comparison is quantified through noise analysis of the images. This system has been implemented for laser scanning microscopy but has potential for widefield fluorescence imaging suitable for open-field surgery.

## 1. Introduction

Light microscopy and optical imaging are among the most important analysis methods in biomedical research and applications. Specimens are analyzed to image such parameters as fluorescence [1, 2], scattering [3–5], refractive index [6, 7], the nonlinear properties of Second Harmonic Generation [8, 9], and Raman Scattering [10, 11]. Techniques include capturing light spectra, fluorescence lifetime [12, 13], polarization [14] with fluorescence microscopy and light backscattering with optical coherence tomography [15–17]. Other light-based techniques involve endoscopy [18–21], computer vision, and pattern recognition-based techniques that work with regular video cameras [22–24], for example, to obtain surface structure information in a surgery setting. Highly light sensitive techniques include methods involving extrinsic or intrinsic fluorescent reporters in microscopy, marking relevant tissue types or biomolecules, and more recently in clinical applications such as the fluorescent marking of tumors for pathology and surgery [25, 26].

All these methods are affected by background light and in a laboratory environment are typically used in a dark room with additional light shielding around the actual instrument. The transition of many imaging techniques outside a controlled lab environment is typically challenged by conflicting lighting requirements between the technology and the requirement of human operators in the setting that typically require a well-lit environment flooded with white light. Common methods to address this problem include the development of special shielding around the imaging system, which is challenging because it provides a complication to the workflow. An alternative approach is to filter unwanted light with frequency filters [27]. This limits the portion of the spectrum accessible for imaging. Spectrally filtering the room illumination to avoid overlap with instrument detection is expensive and can interfere with the color perception of the human viewers working under the filtered light. Other systems aim to pulse excitation light to maximize the fluorescence brightness without overheating the specimen [28]. This method is limited by fluorophore saturation.

The requirement of the imaging system for a completely dark or inadequately lit environment is problematic in many clinical and industrial settings, especially when using robots or other dynamic equipment that interferes with instrument shielding and carries the potential for injury of personnel. Current lighting requirements dictate the layout of laboratory space and workflow, requiring dedicated rooms and causing scheduling conflicts in multi-station labs between instrument setup and image acquisition. Working in darkness can affect safety, efficiency, and causes mental fatigue for lab personnel. Capture requirements also conflict with the need for computer monitors, control lights and illuminated exit signs.

In this paper, we demonstrate a time-based filter to separate one or more instrument detection channels from an ambient room illumination channel. Our transient lighting system exploits the time averaging properties of light detection in the human eye to create illumination that is indistinguishable from normal room or daylight to a human viewer, but leaves most of the available detection time exclusively to optical instrumentation. This is achieved through a light installation that illuminates the room in pulses at a high repetition rate (>100Hz) and a low duty cycle (~1% - 10%) with high brightness, keeping the time-averaged intensity at the level of typical room lighting. While the human eye only perceives the average intensity, detection hardware is gated to capture only in time periods when the room illumination is turned off. We demonstrate this technique on a clinical multiphoton microscope prototype.

## 2. Methods

Our compact automated multiphoton microscope (CAMM) [29], designed to fit on a cart for mobile clinical use, uses a mode-locked Ti:Sapphire laser (Coherent MIRA) as an excitation source, centered at 890 nm with pulse length set to approximately 100 femtoseconds. The excitation light is point scanned by two galvanometer driven mirrors (Cambridge Technology) in a square raster pattern. Depending on capture time and image parameters, the microscope scans a horizontal line in 1 to 15 ms. For this study, a constant horizontal scan time of 1.32 ms was used. Between capturing two lines, no data is captured for 0.62 ms while the galvanometer scanner reverses direction to scan the next line, referred to as the flyback time. A second galvanometer scans vertically across the image over a period of 984 ms, separated by an 8 ms vertical flyback time. Definition of image capture parameters, such as zoom, pixel dwell time, and display of the captured image are handled by a custom lab developed software acquisition platform called WiscScan [29]. A 20 × 0.75 NA air objective lens (Nikon CFI S Fluor 20X) is used to focus the excitation light onto the sample. The laser pulse excites multiphoton fluorescence or second harmonic at 890 nm and the emission light is detected by a photomultiplier tube (PMT) (Hamamatsu 7422 GaAsP). The system is designed for clinical surgical pathology applications and operation under dim room lighting. Under standard experimental conditions, the capture of bright and spectrally narrow multiphoton fluorescence signals is possible but requires the use of a narrow band-pass filter, a shroud around the microscope stage and objective to block room light, and low gain on the PMT. However, multiphoton fluorescence signals tend to be significantly weaker and distributed over a wider bandwidth making them harder to capture. For this paper, we show that high quality multiphoton fluorescence images can be captured in a brightly lit environment, without a protective shroud, by using our transient lighting technique. Without the use of transient lighting, it is only possible to image optimal fluorescence with the room illumination completely turned off.

The imaging process is divided into two or more channels that are separated in time. Each channel receives a certain amount of time in a cycle. After each channels’ time has passed, the next cycle starts. Our cycles consist of a short room illumination channel followed by a much longer fluorescence excitation and capture channel. Switching is coordinated by an Arduino™ microcontroller (SparkFun Electronics, Boulder, CO). We investigate two different setups, both of which are diagrammed in Figure 1.

**Figure 1:**
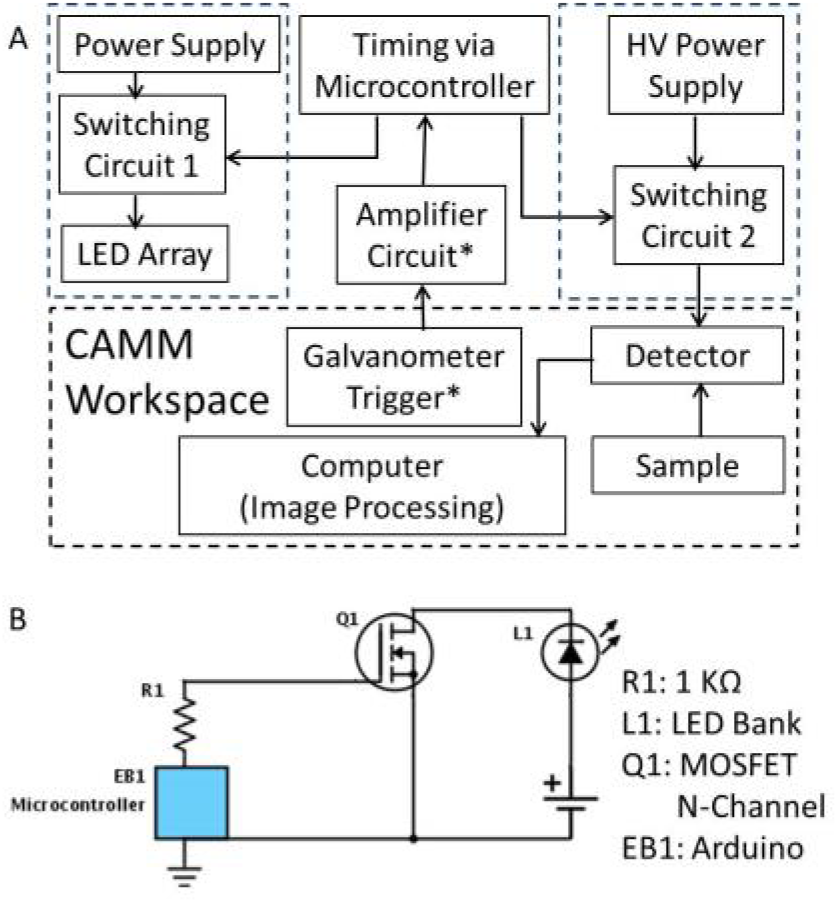
(A) Block diagram of the experimental set up. Boxes with ‘*’ were only used in Setup ‘B’, (B) Circuit diagram for switching via microcontroller using a MOSFET transistor.

### Setup A

The microscope PMT is switched off during the illumination cycle using a high voltage solid-state relay (Panasonic AQV259). The capture process is divided in cycles of 10 ms. In each of these cycles, 1 ms is reserved for illumination, 7 ms are reserved for capture, and 2 ms are reserved as a buffer between channels to allow time for switching. Due to the internal capacitance of the PMT and limited switching time of our solid state relay, the switching process takes about 1 ms. During this switching time, neither illumination nor capture are active.

### Setup B

The PMT is left active and the illumination occurs during the horizontal galvanometer motor flyback time when the microscope is not reading data from the PMT. In this configuration, it is important to protect the PMT against excessive light levels and to prevent light from directly entering the PMT to avoid damage.

For our prototype, we use 64 5W white light emitting diode (LED) lights (Cree Inc., Durham NC) to provide room illumination. The entire set of LEDs can provide 89 lumens (comparable to a bright desk lamp) at 10% duty cycle, providing adequate illumination of the immediate surroundings of the microscope. The light output of the LED is proportional to its power consumption over its operating range. Since the voltage changes only slightly over that range brightness is thus roughly proportional to the current applied to the LED. LEDs and switching electronics are mounted on a custom 3D printed fixture that is inserted in the metal frame of standard desk lamp. Images of the LED fixture and lamp are shown in Figure 2.

**Figure 2:**
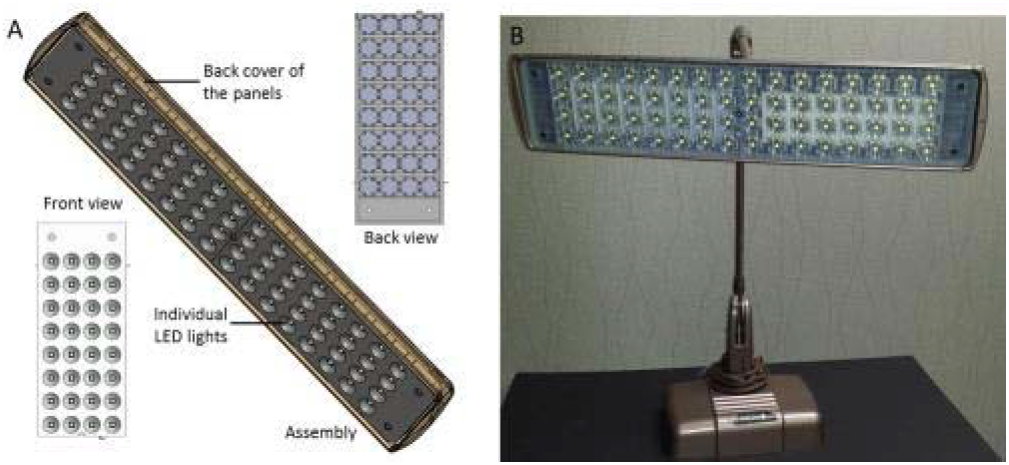
(A) CAD model of the LED array lamp fixture designed in Solidworks™ (Dassault Systèmes, Vélizy, France) showing front and back view of the panel, (B) 3D printed and assembled LED lamp fixture housing 64 LED lights.

In Setup A, we operate the LED light bank at a frequency of 100 Hz with 1 ms on and 9 ms off cycle. In Setup B, the lights are triggered by the horizontal galvanometer drive signal and the cycle frequency is determined by the capture speed of the microscope. The standard image capture rate used for our system, governed by the vertical galvanometer speed, was about 1 frame per second. The trigger signal is detected by a digital input pin of the Arduino™ microcontroller that switches the illumination on for 97 μs every 1.3 ms after the flyback trigger is detected. The speed of switching in the current system is limited by the speed of the microcontroller, which switches between the on state and the off state in about 1 microsecond. The LEDs themselves can switch at faster times, typically about 10 nanoseconds. The system works at any scanning speed as long as the duration or brightness of the lighting pulse is adjusted to maintain a constant average light level. The duration of the lighting pulse is easily customized through serial communication with the microcontroller, but is chosen here to be much shorter than the flyback and hold times when image data is not being collected. The galvanometer signal along with the switching signal applied to the transient lights is illustrated in Figure 3. By triggering in this fashion, the lighting pattern is automatically updated to reflect any changes made in imaging parameters (frame rate, zoom, resolution, etc.) in the WiscScan software.

**Figure 3:**
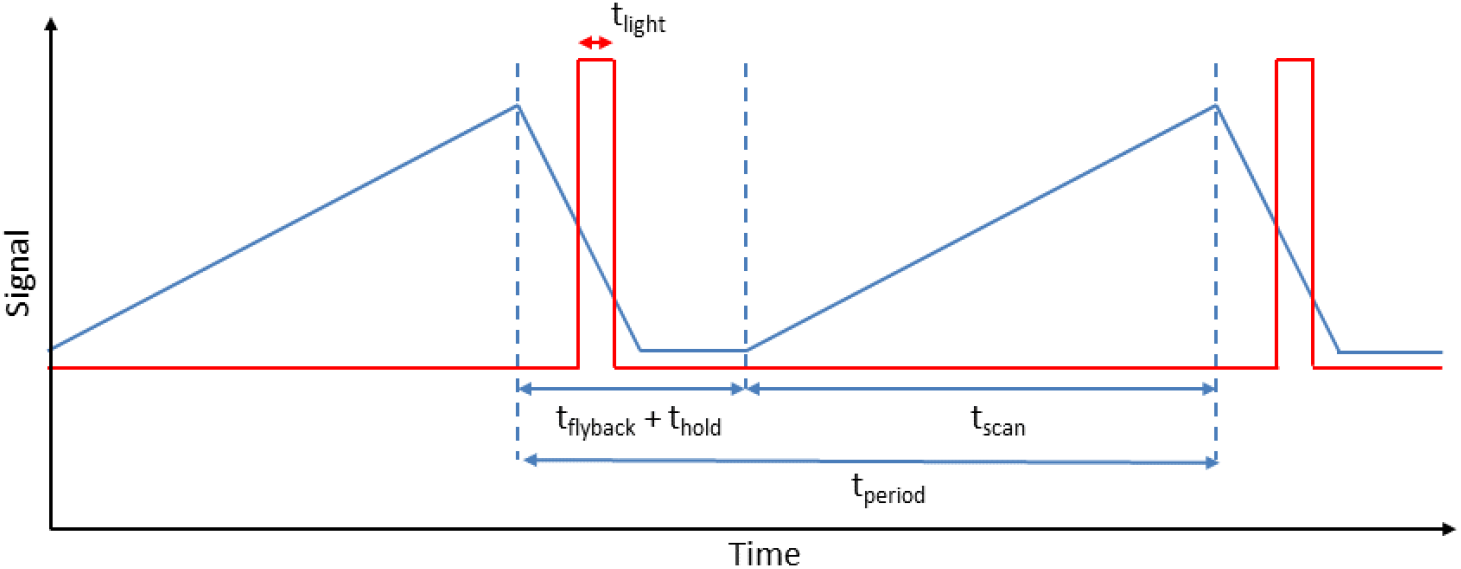
Timing diagram showing the galvanometer feedback signal and the transient lighting switching used in setup 'B'. For the current work, t_flyback_ + t_hold_ is 0.62 ms and t_scan_ is 1.32 ms.

The signal from the PMT is amplified with a C7319 (Hamamatsu, Japan) 200 kHz bandwidth amplifier, and digitized with an A4D4 (Innovative Integration, Simi Valley, CA) data acquisition system controlled by WiscScan. In Setup A, we used a Hamamatsu R374 multialkali PMT. Setup B used a Hamamatsu H7422-40 GaAsP PMT.

In Setup A, the captured images contain two distinct artifacts due to the lighting scheme. First, the image contains patterns of blank lines corresponding to the period when the voltage to the PMT was turned off. Second, periodic spikes are observed at the end of each blank period, corresponding to a signal spike, likely due to accumulated space charge that occurs while the PMT is turned off. These spikes are followed by a period of reduced PMT sensitivity. The background signal of the PMT after reactivation appears at white lines in Figure 4A. To provide a clean microscope image, the data corresponding to these blank lines and spikes need to be discarded and filled in from different frames. This is done via post-processing in MATLAB™ (MathWorks, Natick, MA), in which the invalid pixels in each captured image are identified and a single image containing only valid pixels is composed from several images of the sample. In Setup B, since the illumination entirely occurs during the flyback time and the PMT is not switched, the artifacts apparent in Setup A are non-existent. The captured images thus require no post-processing.

**Figure 4:**
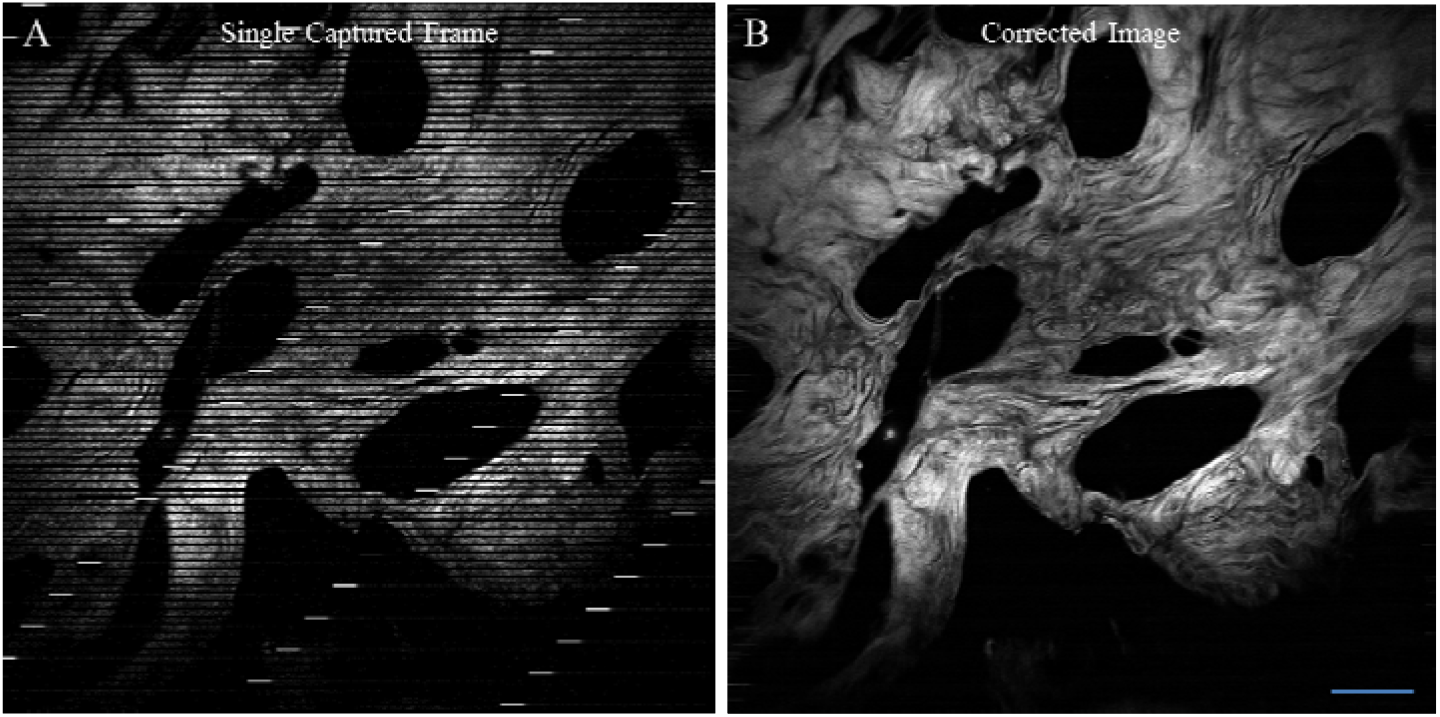
Image captured in setup A with the PMT (Hamamatsu R374) switched off during the illumination cycle. Illumination is not synchronized with the microscope galvanometers in this setup and illumination cycles appear as black segments in the frames (A). 25 frames are combined to generate a complete image (B). Scale bar = 100 μm.

For setup A, a sample of normal human breast tissue was obtained through the University of Wisconsin Biobank program. This tissue was paraffin embedded, sectioned at 5 μm thickness and stained with hematoxylin and eosin. We imaged two photon excited eosin fluorescence in this sample using an excitation wavelength of 890 nm and an emission filter centered at 520 nm with a 30 nm bandwidth. For setup B, we imaged a commercial slide of bovine pulmonary artery endothelial cells (BPAEC) (Life Technologies, Carlsbad, CA). The mitochondria in the live BPAEC were stained by MitoTracker^®^ Red CMXRos with accumulation dependent upon membrane potential. Filamentous actin was labeled with Alexa Fluor^®^ 488 phalloidin while nuclei were counterstained with the blue-fluorescent DNA stain DAPI. The sample was excited at 890 nm The fluorescence was imaged using both a 520 nm filter to capture the fluorescence of the filaments and 492 nm short pass filter to capture fluorescence of the nuclei. The mitochondria were not imaged, as the peak of the excitation spectra for MitoTracker^®^ Red CMXRos was far from the wavelength of the excitation source. Figure 5 shows a transient lighting image of the BPAEC slide where images using both filters are merged to simultaneously visualize actin filaments and nuclei.

**Figure 5:**
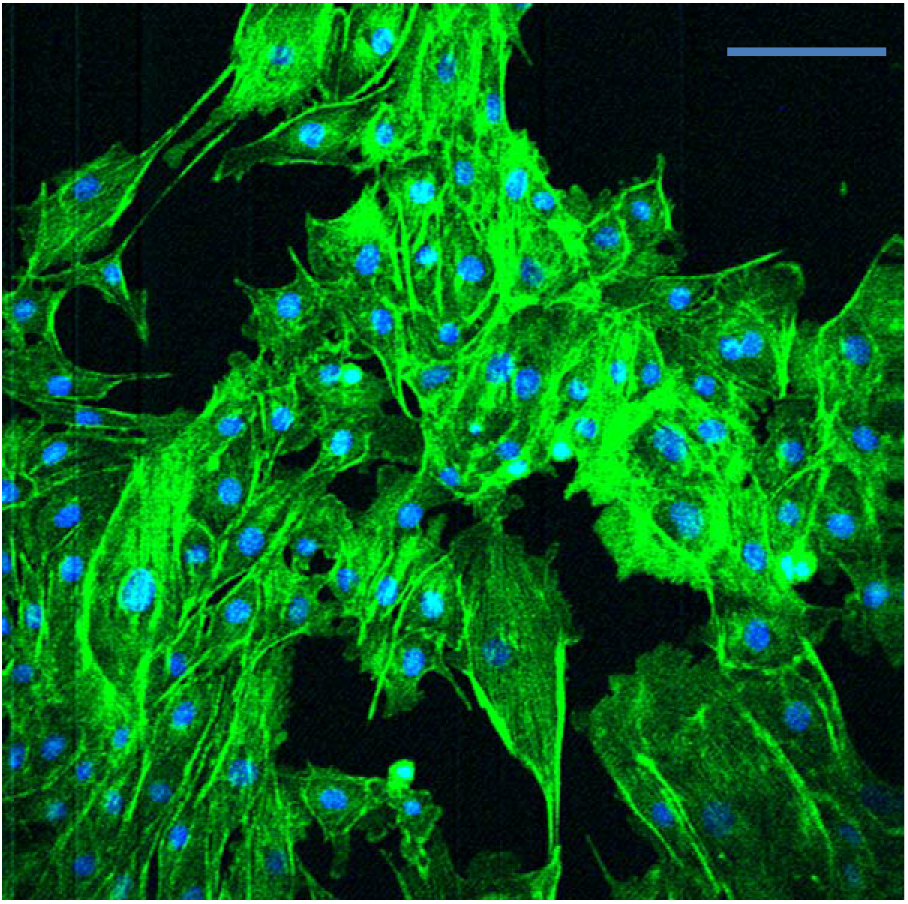
Image of F-actin (green) and nuclei (blue) in BPAEC cells captured in setup B. Illumination is synchronized with the microscope galvanometers in this configuration. Individual images with 492 nm shortpass and 520 nm filters were combined to generate this image. Scale bar = 100 μm.

## 3. Results

First, we characterized the brightness of our transient lighting LED array in comparison to the light provided by the standard fluorescent room lights in our lab. Then, we demonstrated the use of our transient lighting system in both setup A and setup B. The results of these tests are given in the sections below.

The brightness of the transient illumination is measured in several positions around the microscope for normal room light and transient light in both capture configurations. The results are listed in Table 1 and show that the light levels provided by our transient lighting system on average were greater than or equal to the full light levels provided by the standard room lights in the work area around our microscope during our imaging experiments. The results of our imaging experiments are given below.

**Table 1:** Ambient light levels at different positions near the experimental setup for different lighting scenarios.

| Lighting Condition | Light Measurement (Lux) at sample stage | Light Measurement (Lux) at 1m from light source |
| --- | --- | --- |
| No room lights | 12* | -- |
| Room (fluorescent lighting) | 130 | 1000 |
| Transient lighting with LED** | 414 | 800 |

\*Includes light from computer monitor
\*\*Measured at 5% duty cycle

As a proof of principle for our technique, we established experimental setup A, which interleaved the PMT power with illumination and had no actual connection with our microscope scanning system. Figure 4A shows a single frame captured with this setup. Dark lines in the image correspond to illumination cycles when the PMT is switched off. After the PMT is switched back on, a bright signal is detected that saturates the PMT. Multiple frames captured in this way are combined to create one complete image of the sample in Figure 4B. Position of the transient lights and the presence of a protective shroud around the PMT do not affect the image or the PMT signal, which is further described in the discussion.

Setup B integrated our prototype into the scanning system of the microscope to allow for uninterrupted imaging. Images and lighting levels captured with setup B are shown in Figure 6. The top row of panels shows microscope images of the sample with identical settings but under different room lighting conditions. The bottom row shows a photograph of the sample stage under the same lighting conditions. To allow a comparison of lighting levels, all photographs were taken with the ISO800 exposure settings without using a tripod on a Samsung Galaxy S4 camera phone. In Figure 6A and 6D, normal room light is on. The sample and microscope environment are well lit, but the PMT detector is completely saturated, indicated by the red color in Figure 6A. Turning off the room lights enables capture of the sample with the PMT, but the ambient light is insufficient to create a photograph (Figure 6B and 6E). Transient lighting, on the other hand, creates a well-lit scene (Figure 6F) but does not notably affect the microscope image (Figure 6C).

**Figure 6:**
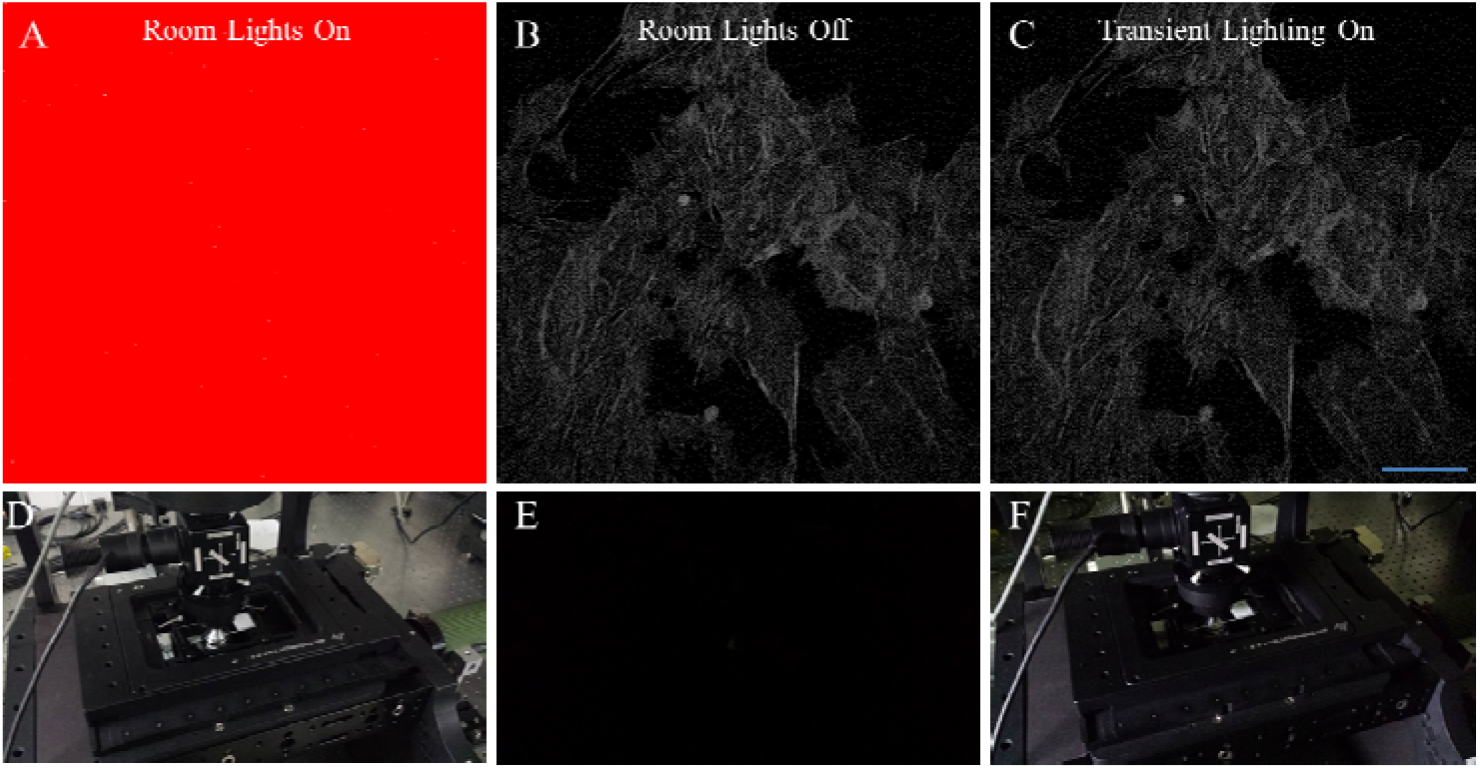
Multiphoton microscope images captured by our system in setup B with the room lights on (A), room lights off (B) and the transient lighting system on with room lights off (C). The imaging parameters were held constant for all images. The photographs in the second row show the microscope stage and sample with the room lights on (D), room lights off (E), and transient lighting system on (F). This figure demonstrates that the transient lighting system can fully illuminate the workspace, equivalent or better than the full room lights, with no effect on the microscope image quality. Scale bar = 100 μm.

## 4. Discussion

When switching off the main power supply of the PMT, the accumulated charge on the PMT dynodes leaves via resistors. The actual voltage drop after switching the PMT off is thus an exponential decay if no photocurrent provides additional discharge. A switching time below 1 ms was observed experimentally and is in agreement with the specifications of the high voltage relay and the discharge times expected for the PMT dynodes. After switching the PMT back on, the dynodes recharge with the same time constant. Immediately after the voltage is switched back on, a peak in PMT current is detected. We attribute this peak to space charges from electrons generated at the PMT photocathode while the PMT voltage is switched off. The PMT signal was studied with an oscilloscope to confirm that this spike is not high enough to damage the readout electronics at any PMT driving voltage. Following this peak is a period during which the PMT generates an elevated background current. The duration of this period is dependent on PMT voltage and lasts less than 1 ms.

Figure 7 shows the transient signal of the PMT when switching on the high voltage with the solid state relay, while looking at only the dark room background light. The plot is generated from data captured by the CAMM. At a typical capture voltage of 600 V (Figure 7A), the signal returns to a normal background after about 600 μs after switching. At 800 V (Figure 7B), this time is doubled. To overcome some of these challenges in our next prototype iteration, we plan to switch the PMT dynode potentials individually as described in our photon counting discussion below.

**Figure 7:**
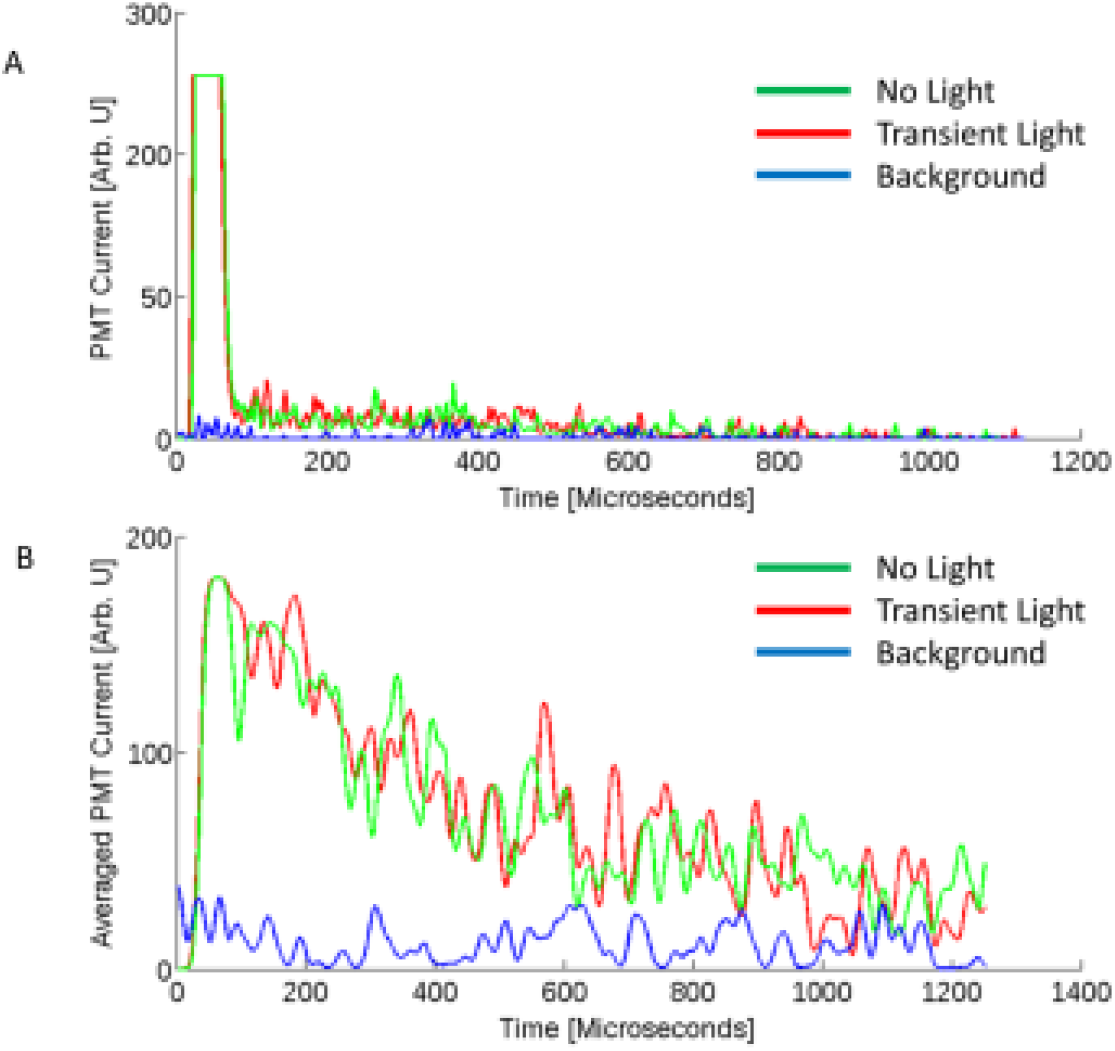
PMT Recovery after switching at (A) 600 V and (B) 800 V. The PMT (Hamamatsu R374) recovers to full sensitivity after less than 1 ms. The initial spike is possibly due to space charges. It saturates the readout amplifier but is well below its damage threshold.

Most commercial lighting LEDs are blue or ultraviolet diodes that contain a fluorescent material that converts the diode light to broadband white illumination. Both the diodes and common fluorescent materials have fluorescence lifetimes in the nanosecond regime and can be rapidly switched on and off.

The green plots in Figure 7 show the PMT signal after switching in a dark room, while the red plots show the switching response with active transient lighting. In this configuration (Setup A), the PMT was exposed to bright LED light just before switching. There is no apparent effect of the LED light on the PMT in this measurement setup.

Leakage from the LED into the PMT cycle can be detected when placing an LED directly on the aperture of the PMT and observing the PMT signal with an oscilloscope. In this case, a signal clearly originating from the LED is detected by the PMT that lasts about 1 ms into the capture cycle. This “afterglow” of the LEDs may be present due to a combination of effects including thermal radiation or particularly long lived fluorescent states in one of the LED materials or any other materials present in the setup. It could also be due to space charges accumulating in the LED. From our analysis we can rule out this afterglow as a contribution to noise in our images (in Setup B). Under normal experimental conditions, the PMT is never exposed to LED light of comparable strength since protected by a filter and directed towards the sample.

Figure 8 and 9 compares the noise in images collected with and without transient lighting and using a 520/30 nm and a 492 nm shortpass filter respectively. The images in Figures 8A and 9A are collected in a dark room, Figures 8B and 9B are collected with active transient lighting in setup B. Histograms of the areas marked in the images are provided in Figures 8C and 9C. To visualize the noise in both images, 10 images were captured with and without active transient lighting and the mean and standard deviation were computed for each image pixel. The standard deviation divided by the mean is shown for each pixel in Figures 8D and 8E for 520 nm filter; and Figures 9D and 9E for 492 nm filter. A histogram of the indicated region in the image is shown in Figure 8F and 9F for respective filters. Our analysis does not reveal a notable difference in noise between the images created with and without transient lighting.

**Figure 8.**
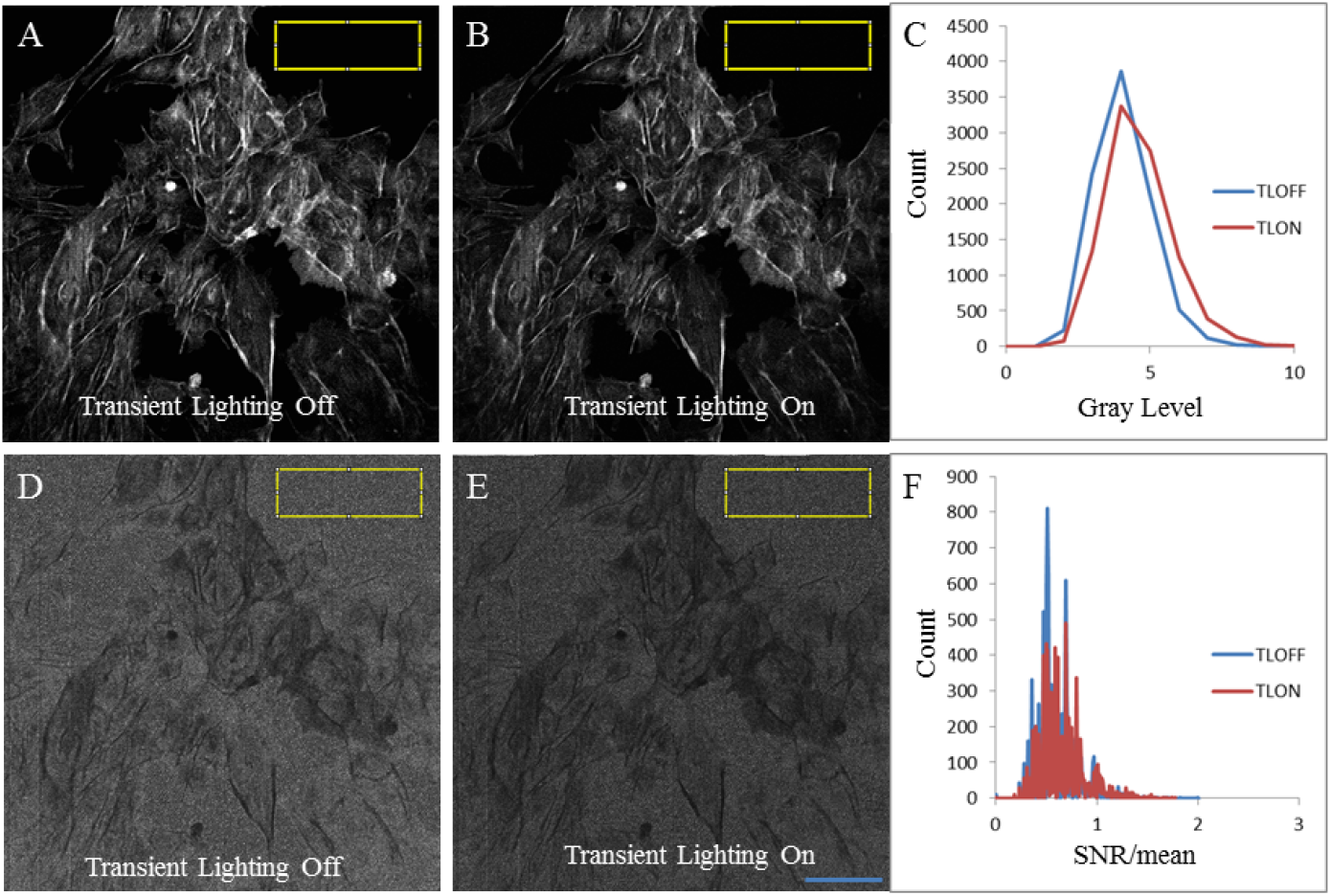
Average of ten microscope images with the room lights off (A) and transient lighting (TL) on (B) using the 520 nm filter. Histograms of gray levels within the yellow ROI for the room lights off (blue) and transient lighting on (red) images (C). Standard deviation normalized by the average gray level for the image with the room lights off (D) compared to the transient lighting on (E) with histograms of gray levels (F) within the yellow ROIs for the room lights off (blue) and transient lighting on (red). Scale bar = 100 μm.

**Figure 9.**
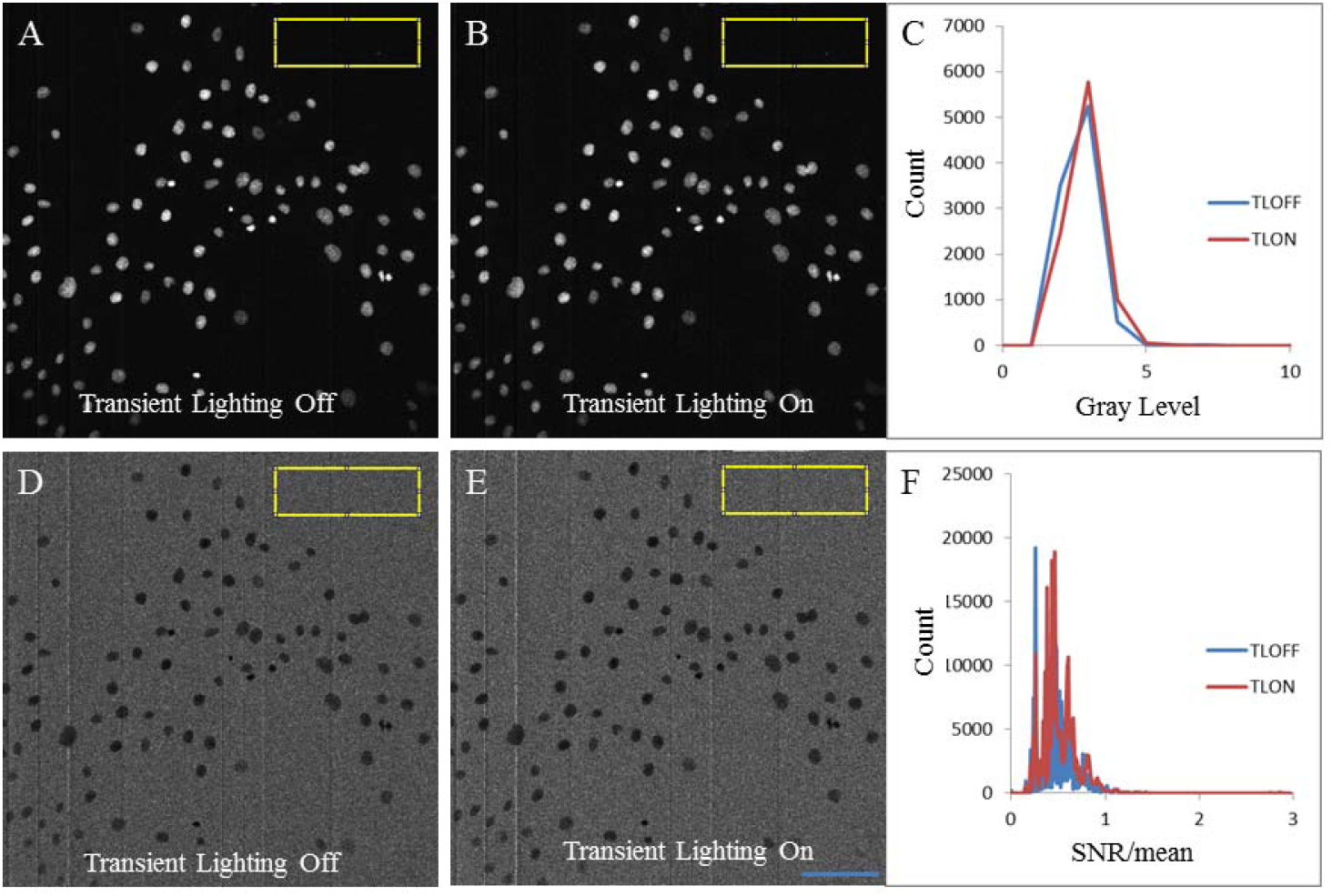
Average of ten microscope images with the room lights off (A) and transient lighting(TL) on (B) using the 492 nm shortpass filter. Histograms of gray levels within the yellow ROI for the room lights off (blue) and transient lighting on (red) images (C). Standard deviation normalized by the average gray level for the image with the room lights off (D) compared to the transient lighting on (E) with histograms of gray levels (F) within the yellow ROIs for the room lights off (blue) and transient lighting on (red). Scale bar = 100 μm.

It has been shown [30, 31] that light intensity modulations above a frequency of 200 Hz are neither recognized by the human brain, nor physically detected by the sensory receptors in the retina. Light at frequencies of about 60 Hz under short duty cycles cause headaches in some individuals in certain situations, while a visible flicker is perceived at 30 Hz. Frequencies at around 13 Hz are known to cause seizures in some individuals. Most traditional artificial illumination is in fact transient. Incandescent light bulbs are modulated at a frequency of 60 Hz but the heat decay constant of the filament provides ample intensity smoothing.

Fluorescent and commercial LED lights use 100 Hz or higher frequencies. Light from pulsed laser sources is equivalent to continuous laser light from an eye safety point of view as reflected in laser safety regulations, as long as the pulse repetition rate is high, and the pulse peak intensity is not large enough to excite nonlinear responses in the eye. Based on all this information we conclude in summary that, there is no known mechanism that could cause any adverse health effects due to a transient lighting illumination scheme if the cycle repetition rate is kept above 300 Hz and room illumination duty cycles are on the order of 1% or higher. Lower repetition rates of about 60 Hz will likely cause headache or migraine symptoms in some individuals under continuous regular exposure (such as 8 hours per day for weeks). 100 Hz flicker frequencies are used in low frequency incandescent lighting and while there are known effects of these systems they are currently widely accepted for lighting [32].

## 5 Future Work

Both PMTs used in our system are suitable for photon counting microscopy setups. Our microscope currently lacks the readout electronics to perform photon counting measurements. Measurements on our system are currently performed with a PMT gain of 10^5^ or less. This is sufficient for many analog microscopy applications. Photon counting is usually performed with a PMT gain of 10^6^ corresponding to a PMT voltage of 1100 V for the R374 PMT. When operated at this voltage, the PMT requires at least several milliseconds of recovery time according to our measurements. We conclude that the configurations presented here are not suitable for a photon counting setup but will be the focus of future work given the advantages of photon counting.

The stability and switching speed of the PMT can be improved by switching the potentials of individual PMT dynodes rather than the entire PMT. Modulation of individual dynodes to modulate PMT gain is routinely performed at nanosecond speeds, but with insufficient modulation depth around 80%. By directly switching multiple dynodes of the PMT, it may be possible to switch a PMT fast enough for photon counting transient lighting.

PMTs are not the only light sensitive detector that can be used with transient lighting. CCD and EMCCD cameras with microsecond speed electronic shutters are commercially available. Placing a shutter in front of the actual sensor is another viable option. Electronic shutters based on liquid crystals are widely used but only transmit one light polarization in their open state and thus contribute additional light loss. Mechanical shutter of sufficient speed could be built from inexpensive parts and has broad spectral bandwidth and a large contrast ration between the open and closed states. Commercially available motorized iris shutters are fast enough to work for our system, but are usually not designed to survive more than a few hundred thousand open and close operations and therefor would wear within a few hours of operation. A more workable solution is a chopper wheel with two blades rotated in front of the detector. For a detector aperture of 1 cm a wheel of about 15 cm diameter rotated at 3000 rpm would be sufficient to achieve a 100 Hz lighting frequency. Because of the choppers inertia it is slow to respond to changes in lighting frequency for example due to a change in frame rate. Mechanical shutters have the advantage that they can be applied not only in microscopes but also in spectrometers, endoscopes or wide field fluorescence monitors. The problem of room light interference can be more severe in a wide field camera than it is in a microscope since a wider field of view collects light from a larger area.

Through the use of a high-speed, triggered camera, it is possible to implement transient lighting in a wide field fluorescence imaging system. Among many applications, wide field fluorescence cameras may be used in hospitals in conjunction with fluorescently labeled tumors [25] or other optical signatures for both pathology and intraoperative use. Shielding ambient light in these applications is difficult, as the body itself is not a good absorber at some wavelengths and thus in a surgical setting some of the background light may emanate as scattered light from the patient and cannot be shielded. Often, the lights are simply turned off when fluorescence guidance is used, creating safety and comfort issues for surgical staff.

Utilizing transient lighting in these applications has the potential to improve the sensitivity and flexibility of these procedures, due to its high ambient light rejection and wavelength independence.

## 6. Conclusion

We have presented a transient lighting system and evaluated its performance using a multiphoton fluorescence microscope system. The multiphoton implementation was an excellent use case due to the sensitivity of this system. By allowing for highly sensitive optical detection to take place in well lit spaces, transient lighting can aid the transition of many promising imaging methods into clinical and industrial applications. It also provides a viable alternative to wavelength filters in situations where broadband spectral information is important. The increased utilization of LED lighting installations makes it feasible to implement this method on a larger scale for entire buildings and cities.

## Acknowledgments

We would like to acknowledge Robert Swader at the Morgridge Institute for Research (MIR), Madison, Wisconsin for help with CAD design and 3D printing of the transient lighting fixture. We would also like to acknowledge Adib Keikhosravi for his help with CAMM at the Laboratory for Optical and Computational Instrumentation (LOCI), at the University of Wisconsin-Madison. We also acknowledge project funding from the MIR and the LOCI.

